# Input Construction Accounts for Most PoseBusters Clash Failures on RNA–Ligand Complexes

**DOI:** 10.64898/2026.11.24.768943

**Authors:** Mahdi Soultana, Olamide Isreal Odesanya, Mervin Gaitho, Gabriel Richman

## Abstract

PoseBusters, a widely used test of the physical validity of ligand poses, was calibrated on protein–ligand crystal structures, and whether its checks transfer to RNA has not been measured. Applied without filtering the input to 2,970 deposited RNA–ligand poses, the receptor-clash check failed 527 of 2,931 (18.0%), and all but 7 of the failures were in covalently bonded nucleotides or in entries that contain protein, were solved by cryo-electron microscopy or were resolved at worse than 2.0 Å, so most failures are associated with which structures and ligand records entered the input set. On a high-resolution set constructed after these failures were attributed, the check failed 3 of 149 (2.01%; 95% confidence interval (CI) 0.00–4.58%) against 0.32% for the protein benchmark, a difference this set cannot resolve, and the PoseBusters calibration rule selected a threshold of 0.735 against the shipped 0.75. We give a seven-step dataset construction protocol and a tool, rnaprep, that implements its first four steps. On 139 redocking targets, with accuracy measured in the crystal frame without superposition, requiring a valid pose cost NucleoDock 25.2 points (69.06% to 43.88% within 2 Å) and AutoDock Vina none, and cut the lead of NucleoDock from 38.9 points (95% CI 26.5–50.0) to 13.7 points, a lead not clearly distinguishable from zero (95% CI 1.3–27.5; cluster permutation P = 0.053). For these two programs in one redocking regime, the result supports reporting validity alongside RMSD in RNA docking benchmarks.

## 1. Introduction

A docked or co-folded ligand pose can lie within 2 Å root-mean-square deviation (RMSD) of the crystallographic answer and still contain steric clashes or distorted geometry. PoseBusters [1] separates the two axes, testing chemical validity, intramolecular geometry and intermolecular contacts. Its thresholds are set so that crystal ligands pass: “All crystal structures in the Astex Diverse set and all but one in the PoseBusters Benchmark set pass at this threshold”. Both reference sets, the Astex Diverse set [2] and the PoseBusters Benchmark set, are protein–ligand complexes.

RNA–ligand docking is evaluated mainly by RMSD, which does not show whether the poses of a top-ranked program are physically plausible. This holds for the dedicated nucleic-acid scoring functions and docking programs [3–6], for the comparative assessments of those programs [7–9], for the pose-reranking and benchmark-set work built on those assessments [10,11], for learned estimators of nucleic-acid pose accuracy [12], for co-folding models that predict nucleic-acid complexes [13–16], and for a recent review of the field [17]. A screen of the works citing PoseBusters and HARIBOSS, a literature search and the PoseBusters issue tracker, all on 13 November 2026 (S1 File, Section S7), found no study that measures how often deposited RNA–ligand structures fail PoseBusters checks, or that explains why they fail (queries and sources are listed in that section; the search covers the indexed literature of that date only).

One study found in that screen reports PoseBusters pass rates per docking program. NucleoDock [18] states that “PoseBusters criteria that are evidently incompatible with the energy and chemical characteristics of nucleic acid-ligand systems were excluded … Without this adjustment, all docking methods would yield zero success rates”, and gives as an example that “certain ligands in nucleic acid complexes intrinsically contain radical species, a feature that is also observed in experimentally resolved complexes deposited in the PDB”. Which checks were excluded is not stated. Its supplementary tables still report the receptor-distance and volume-overlap checks, and its main text advises that “these receptor-contact checks should be interpreted conservatively in the nucleic acid–ligand setting”. It reports pass rates for docked poses only. Whether the protein-calibrated thresholds hold on deposited RNA–ligand crystal structures has therefore not, to our knowledge, been measured.

The aims of this study were to (i) measure the rate at which deposited RNA–ligand poses fail each protein-calibrated PoseBusters check and find what drives it, (ii) derive nucleic-acid thresholds from those crystal structures by the same calibration rule PoseBusters applied to protein complexes, and (iii) determine how far a reported docking success rate moves when a pose is required to be physically valid as well as within 2 Å RMSD, for a general-purpose and a nucleic-acid-specific docking program on the same targets. Because the PoseBusters thresholds are set so that crystal ligands pass, high-resolution crystal structures are expected to pass. The result that carries the paper is the first: most of the failures under naive application came from how the input set was built, and on a correctly constructed set of high-resolution crystal structures the receptor-clash check failed at a rate that could not be distinguished from the protein reference. For the second, the calibration rule selected a receptor-clash threshold of 0.735 against the shipped 0.75, so no new threshold is proposed. For the third, requiring validity cut the RMSD lead of the nucleic-acid-specific program over the general-purpose one to about a third, leaving a lead these data do not clearly distinguish from zero, in a two-program test that shows the size of the effect and does not rank programs.

## 2. Materials and methods

### 2.1 Data sources

Four sources were combined. An RNA-only set was built from the RCSB Protein Data Bank on 12 November 2026 by searching for entries with at least one RNA chain, zero protein chains and at least one non-polymer entity; DNA-containing entries are excluded later, where “RNA-only” is defined (Section 2.5). HARIBOSS [19] contributed 2,131 entries through its CSV export. The 173-structure benchmark set of Agarwal et al. [7] was taken from Table S1 of that paper, and the 125-complex set of NucleoDock [18] from its Table S2, which that study assembled from the four benchmark sets of Yan and Wang, Ruiz-Carmona et al., Chen et al. and Philips et al. [5,20–22]. The union contained 2,244 entries, of which four were obsolete in the RCSB Protein Data Bank and were dropped.

### 2.2 Dataset construction

A chemical component was retained if HARIBOSS listed it for that entry, or if it passed an additive filter: not a crystallization additive, ion or solvent; at least 10 heavy atoms; molecular weight between 100 and 900 Da. One instance was taken per entry and component, the instance HARIBOSS names where it names one. In 11 of the 2,240 current entries no component passed, because each held only components that HARIBOSS did not list and the additive filter excluded, or no non-polymer component at all. The final set contained 2,970 ligand instances over 2,229 entries.

Each instance was resolved by chemical component identifier, `auth_asym_id` and `auth_seq_id`. Covalent linkage to the polymer was recorded two independent ways: a deposited `struct_conn` record of type `covale` joining the instance to a polymer residue, and the shortest heavy-atom distance from the ligand to any polymer atom.

### 2.3 Structure preparation

For each instance, the 10 Å pocket was retrieved from the RCSB ModelServer as mmCIF with all categories, which carries the deposited `struct_conn` records, and converted to PDB format with gemmi 0.7.5. The ligand was retrieved as SDF for that instance alone. The receptor was defined as the pocket minus every non-polymer copy of the ligand component, including copies other than the scored instance, and minus water, matching the PoseBusters Benchmark construction (“PyMOL was used to remove solvents and all occurrences of the ligand of interest”).

Pocket and full-receptor preparation gave identical results for every check on 373 of 376 instances. In the other three the full-receptor query lacked the residue number and scored a different copy of the ligand, and only the receptor-clash check differed.

Seven ligand instances, of the components C, G and N, share their identifier with a polymer residue name. For these, the ligand was removed from non-polymer residues only, so that polymer nucleotides of the same name stayed in the receptor. One of the seven, 8P2K/N, has no polymer atom within its 10 Å pocket, and PoseBusters raised an exception on the empty receptor. It is the one instance of 2,970 that was not checked.

Each receptor kept only atoms with a blank alternate location or the ligand’s own. Of the 2,970 ligands, 56 sat in a non-blank alternate location, and 35 of their receptors also held atoms in another one, which were removed. This changed the receptor-clash outcome of three instances (6C8N/EQ1, 5SWD/ ADE and 6P1U/CTP) and the organic-cofactor outcome of four, all from fail to pass. The rule assumes that the deposited alternate-location labels are consistent within the pocket. Removing receptor atoms can only turn failures of the minimum-distance clash check into passes with all other inputs fixed, and where the labels are inconsistent it can hide a real contact (S1 File, Section S1).

### 2.4 Software and parameters

PoseBusters 0.6.5 was run in the `dock` configuration, unmodified and at its default thresholds, with the full report enabled. Redocking used AutoDock Vina 1.2.7 [23,24] with ligand preparation by Meeko 0.8.0, a search box centered on the deposited ligand with 5 Å padding, exhaustiveness 32 and seed 12648430, matching the protocol of the source benchmark [7]. The deposited ligand was read from its SDF file with hydrogens added. The receptor was typed for Vina from the element column of the PDB file (AutoDock atom types, all partial charges zero), which does not affect the type-based Vina scoring function. Accuracy was measured as the symmetry-corrected heavy-atom RMSD of RDKit 2026.9.2 (`rdMolAlign.CalcRMS`), computed the same way for both programs and without superposition: the pose and the deposited ligand are both in the receptor’s coordinate frame, so the RMSD measures where the ligand lies in the pocket and not only how well its shape matches. `CalcRMS` takes the best atom mapping over the symmetry of the molecule but moves neither molecule. For discussion, the table of poses also carries the RMSD after optimal superposition (`rdMolAlign.GetBestRMS`) under a separate name (`rmsd_aligned_top1`); no rate in this paper uses it. If `CalcRMS` fails for a pose (atom-order or substructure mismatch), the target has no measurable RMSD, is counted as a failure and is reported; there is no fallback to the aligned value (Section 3.6). The deposited ligand for the rescoring was read from the `lig_dep.pdbqt` that Meeko wrote from the deposited coordinates, which are in the receptor frame, and the pose from the first model of the docking output.

The NucleoDock docking, the PoseBusters reproduction and the `rnaprep` preparation runs were executed on the Omic research platform (https:// omic.ai). An AI coding assistant (Anthropic Claude) was used to write analysis code and draft text.

NucleoDock [18] was run unmodified at its default parameters from its public release (https://github.com/ShiYue3384/NucleoDock, commit `0b931cd2`), driven by a Nextflow wrapper (commit `da92d67f`) that adds no docking logic, on the Omic platform under a GPU profile with Nextflow 26.10.2, one run per target, from a container pinned by digest (sha256:dea0dd003e8d35b1dae2be5c4738b3d6b908978f1e5167951450acb2d2c2dabc) that holds the checkpoint graph_99.ckpt together with the Nucleotide Transformer v2 250M and RNA-FM models, with the seed fixed at 42 by the shipped configuration and one thread in the pose-generation stage. The cluster GPU (compute capability 12.0) is newer than the CUDA 11.8 build of PyTorch in the image supports, so the pose-generation stage ran on the CPU. Each run received the same prepared receptor as the AutoDock Vina run and the same deposited ligand as the box-defining reference. The two programs received the ligand differently: AutoDock Vina started from the deposited three-dimensional ligand prepared by Meeko, whereas NucleoDock, which takes SMILES as input, generated the ligand geometry itself. NucleoDock writes ten poses per ligand; the first was scored, because the rates it reports are top-1. NucleoDock poses were scored with the same unmodified PoseBusters `dock` configuration and the same symmetry-corrected RMSD as the AutoDock Vina poses (`score_nucleodock.py`).

PoseBusters 0.6.5 assigns receptor atoms to its checks by PDB record type. Atoms in ATOM records are scored by the receptor-clash check (`minimum_distance_to_protein`), and atoms in HETATM records other than water and inorganic cofactors are scored by the organic-cofactor check (`minimum_distance_to_organic_cofactors`). Both checks use a van der Waals scale of 1.0 and a clash cutoff of 0.75. Version 0.6.5 contains the fix for PoseBusters issue #96, closed on 9 February 2026; before that fix, organic cofactors were counted as receptor atoms when PDB residue information was present.

### 2.5 Evaluation metrics

The receptor-clash check (`minimum_distance_to_protein`) reports the minimum, over ligand–receptor atom pairs, of the interatomic distance divided by the sum of the two van der Waals radii. A pose fails when that fraction falls below the threshold, which is 0.75 by default. A pose was counted as accurate when its symmetry-corrected RMSD to the deposited ligand, measured in the crystal frame without superposition, was below 2.0 Å, and as valid when it passed all applicable PoseBusters checks. Targets that failed to dock were counted as failures.

The categories below overlap, and a category is a property of the entry rather than a demonstrated cause of the clash. Each receptor-clash failure was assigned to the first applicable category in a fixed order: (i) the ligand carries a deposited covalent bond to the polymer; (ii) the entry contains a protein chain; (iii) the experimental method is electron microscopy; (iv) the X-ray resolution exceeds 3.0 Å; (v) the X-ray resolution lies between 2.0 and 3.0 Å. Failures matching none of these (solution NMR, and X-ray at ≤ 2.0 Å) were reported as residual. “RNA-only” denotes entries with no protein and no DNA chain; the correctly constructed set of Section 3.3 also admits RNA–DNA entries. In the attribution, categories (iv) and (v) are resolution > 3.0 Å and 2.0 Å < resolution ≤ 3.0 Å. In the resolution bins of Section 3.5 and Fig 3, bins are closed on the left, and the 2.5–3.0 Å bin spans 2.5 ≤ resolution < 3.01 Å, so it holds the three instances deposited at 3.000–3.009 Å.

Thresholds were derived from the relative distances of the clean set by three rules: the rule PoseBusters applied, under which all crystal ligands but one pass (the second-smallest value); a protein-matched threshold, the 1/308 quantile, corresponding to the 0.32% false-failure rate PoseBusters reports for its own protein benchmark; and a 95% rule, the 0.05 quantile. Quantiles use linear interpolation between order statistics (NumPy default), and a pose fails when its value lies strictly below the threshold. Because 149 values cannot produce a rate of exactly 0.32%, the number of failures each threshold produces is reported with it.

### 2.6 Statistical analysis

No prospective sample-size calculation was made: the set comprises every instance returned by the sources and filters of Sections 2.1 and 2.2 on the query date, and the 139 docking targets are the complete X-ray, RNA-only, non-covalent subset of the 173-structure benchmark [7]. Every rate carries a 95% confidence interval (CI) unless stated otherwise.

Intervals are percentile intervals from a cluster bootstrap [25] over HARIBOSS sequence and pocket equivalence classes (4,000 resamples, seed 20261113), because RNA structures are highly redundant and binding sites within one family are not independent. The classes are used as HARIBOSS gives them, and an instance whose entry carries no class formed a class of its own. Of the 139 docking targets, 24 carry no class and 48 of the 71 resulting classes hold a single target, the largest holding 19. The unit of resampling is the class in every interval. The all-but-one threshold was bootstrapped the same way (2,000 resamples), each resample recomputing the second-smallest relative distance, so its bounds are observed values.

The two programs were compared on the same 139 targets. McNemar’s exact test is reported with the discordant counts, but it treats targets as independent. Inference therefore rests on a paired cluster bootstrap (10,000 resamples) that resamples whole classes with both programs and all criteria of a target kept together, giving intervals for the difference between programs under each criterion, for each program’s validity cost and for the change in the difference when validity is required, and on a cluster-level sign-flip permutation test for each difference (`paired_cluster_bootstrap.py`). Treating the 24 targets with no class as one cluster instead of singletons left the point estimates unchanged: the interval for the RMSD difference became 27.46 to 51.28 (from 26.61 to 50.30 in the same resampling), and that of the validity-qualified difference at 0.75, which excludes zero with singletons (0.88 to 27.62), included it (−0.79 to 27.12). Receptor-clash failure was regressed on entry properties by logistic regression with standard errors clustered by class [26], with resolution centered at 2.0 Å and filled with the median for solution-NMR entries. The resolution trend was tested by a logistic trend with clustered errors and by a rank correlation with a P value from 10,000 permutations of whole classes (`review_gaps.py`). The interval of the protein reference rate (1 of 308) is the Wilson score interval; that rate comes from the hand-curated protein benchmark of PoseBusters, which differs in curation, ligand selection and software version, so the comparison is indicative. The clean-set rate was compared with it by two-sided Fisher’s exact test, which treats instances as independent and is reported beside the cluster interval.

No analysis was pre-registered and no correction for multiple comparisons was applied. Many resolution cut-offs, calibration rules, strata, docking arms and subsets were examined, and several P values lie near 0.05 (0.053 for the validity-qualified lead; three others between 0.02 and 0.05); all are reported unadjusted and should be read as descriptive rather than confirmatory.

Analyses used Python 3.12 with pandas 2.2.3, NumPy 2.2.6, SciPy 1.17.1 and statsmodels 0.14.6. `headline_numbers.py` recomputes 130 numbers quoted in Results and Discussion from the released tables, `methods_numbers.py` the counts of Sections 2.1 and 2.3, and `gap_numbers.py` the 225 numbers of Sections 2.6, 3.2, 3.3, 3.5 and 3.7 to 3.10 that come from the additional analyses; all three fail if any value differs from the one reported here. Numbers covered by none of them, such as the citation counts of S1 File, Section S7 and the 34.8 Å example of Section 4.4, come from logged sweeps and runs.

### 2.7 Additional docking arms and the overlap check

Four further analyses test the effect of unequal ligand input, of a single comparator program, of a single run and of possible overlap between the 139 targets and the NucleoDock development data, using the same receptors and search boxes as the first two arms. In the matched-input arm, AutoDock Vina 1.2.7 docked an RDKit conformer of the SMILES string that NucleoDock received (ETKDG version 3 embedding [27], seed 42, MMFF94 minimization, Meeko 0.8.0), with the exhaustiveness (32), number of poses (20), seed (12648430), box and receptor preparation of Section 2.4. In the third-program arm, smina 2017.11.9 [28] docked the deposited ligand prepared as for the first arm, with the same box, exhaustiveness, poses and seed, in the container image of the registered smina tool (quay.io/biocontainers/ smina:2017.11.9--0). The AutoDock Vina arm reported here was docked with the protocol of Section 2.4 from the prepared inputs (`run_gap_dock_local.py`), and PoseBusters was run on the receptor rebuilt from the prepared PDBQT, which reproduced all 714 PoseBusters values of six matched-input poses (`score_arm_from_pdbqt.py`). All arms were scored as in Section 2.4 from the first pose, with the unaligned RMSD, with the intervals of Table 4 and the paired cluster bootstrap of Section 2.6 (`gap_arms_analysis.py`; S1 File, Tables S12 and S13). The NucleoDock arm was run a second time on the Omic platform with the same container digest, configuration and inputs, as one batch workflow in place of 139 separate submissions. The 139 targets were compared by PDB entry identifier with the 125-complex external test set of the NucleoDock preprint [18] and with the lists of its public repository at commit `0b931cd2`, and the paired comparison was repeated inside and outside that test set. The unaligned RMSD of the first NucleoDock run, whose poses were not archived, was read from the poses of this repeat, except for 2GDI, 2HOJ and 2HOM, the targets nearest the 2 Å cut, whose first-run poses were read back from the platform output; the AutoDock Vina arm of the first run was regenerated as stated above (95 poses within 2 Å after superposition against 96 in the first run).

### 2.8 Use of AI tools

The work was carried out on the Omic research platform (https://omic.ai), which ran the docking, validity-checking and structure-preparation workflows, stored their inputs and outputs, and recorded every run. The repeat of the NucleoDock run (Section 3.10, 139 targets) was launched on the platform through its agent interface, whose analysis agents use Google Gemini 3.1 Flash Lite by default; its poses were scored with the archived scripts, as for every other arm. Anthropic Claude (through Claude Code, with the Claude Opus and Sonnet models) was used to write and run the analysis, scoring and plotting code, to prepare the platform workflows, to search the literature, and to draft and revise the text of every section. Draft versions were reviewed by large language models prompted as journal reviewers: Claude Sonnet and Opus, OpenAI GPT models (including through Codex), Google Gemini, xAI Grok, DeepSeek and Perplexity Sonar; the authors assessed their comments and decided which changes to make. The authors directed the study, reviewed the code and the text, and checked every number quoted in the text against the archived tables with scripts that fail on any difference (`headline_numbers.py`, `methods_numbers.py` and `gap_numbers.py`; Data availability). Every reference was resolved against Crossref or arXiv. No figure was generated or altered by a generative model; the figures are drawn by plotting code from the archived tables. The interpretations and conclusions are the authors’ own, and the authors take full responsibility for the content.

## 3. Results

### 3.1 Failure rate under naive application

Of the 2,970 ligand instances, 2,969 were successfully checked and 2,931 carried the receptor-clash metric. The receptor-clash check failed 527 of those 2,931 instances (18.0%). The remaining checks failed at rates near zero: the radical check failed 0 of 2,969, sanitization 0.37%, bond lengths 0.37% and the internal steric clash check 0.71%.

### 3.2 Attribution of receptor-clash failures

Each of the 527 failures was assigned to the first category it matched (Table 1, Fig 1). A category is a property of the entry and does not establish the cause of an individual clash.

**Fig 1.**
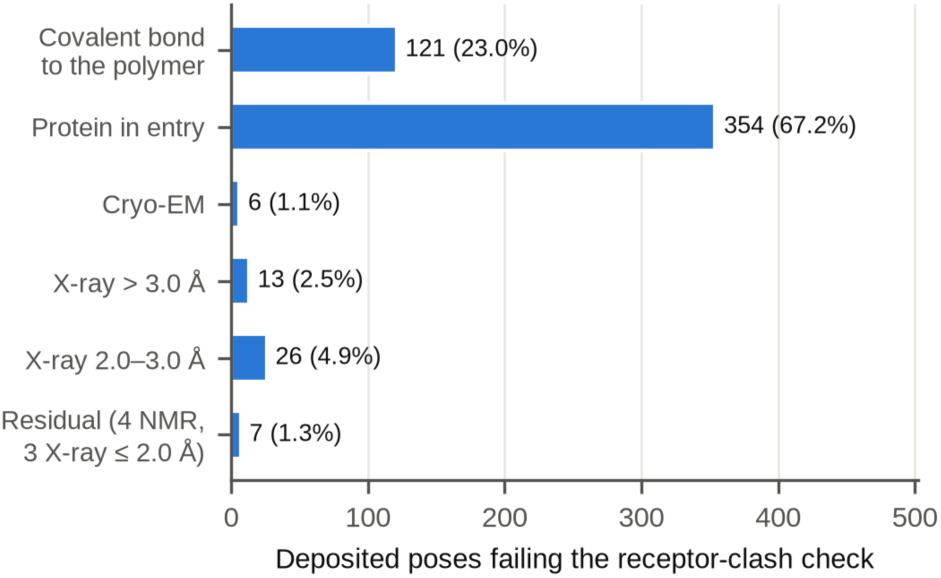
Attribution of receptor-clash failures among deposited RNA–ligand poses. The 527 of 2,931 instances failing the receptor-clash check (threshold 0.75), each assigned to the first applicable cause, tested from top to bottom. Bars are labeled with the count and the percentage of all 527 failures. The residual class holds four solution-NMR entries and three X-ray entries at ≤ 2.0 Å. Cryo-EM, cryo-electron microscopy; NMR, nuclear magnetic resonance.

**Table 1.** Attribution of the 527 receptor-clash failures, in the order the categories were tested. A category is a property of the entry, not a demonstrated cause of the clash.

| **category** | **failures** | **share** |
|---|---|---|
| deposited covalent bond to the polymer | 121 | 23.0% |
| entry also contains protein | 354 | 67.2% |
| electron microscopy | 6 | 1.1% |
| X-ray resolution > 3.0 Å | 13 | 2.5% |
| X-ray resolution 2.0–3.0 Å | 26 | 4.9% |
| residual | 7 | 1.3% |

The categories overlap, so the shares depend on the order in which they are tested. A logistic regression of receptor-clash failure on the same entry properties does not depend on that order (2,931 instances, 527 failures, 900 equivalence classes; standard errors clustered by class; S1 File, Table S9). A deposited covalent record carried an odds ratio of 27.98 (95% CI 16.80–46.61), each ångström of poorer resolution 1.90 (1.50–2.42), protein in the entry 1.51 (0.99–2.31; P = 0.054), DNA in the entry 1.40 (0.87–2.26; P = 0.17), electron microscopy 1.13 (0.73–1.73; P = 0.59) and solution NMR 0.09 (0.02–0.39; P = 0.002). Once resolution is held, electron microscopy shows no detectable association with failure, which the ordered attribution of Table 1 does not show. Of the 354 failures attributed to entries containing protein, 223 were cryo-electron microscopy structures and 89 X-ray structures above 3.0 Å. Under an order that tests method and resolution before protein, only 6 failures are attributable to protein in the entry alone (S1 File, Section S2). The covalent share and the 7 residual failures do not depend on the order.

The seven residual failures were four solution-NMR entries, two X-ray structures at ≤ 2.0 Å with no protein in the entry and one RNA–DNA hybrid at 1.60 Å. Covalent annotation and interatomic distance did not agree, so neither can substitute for the other: 87 instances had a polymer contact below 1.8 Å with no deposited `covale` record, and 45 carried a record without a short contact (S1 File, Section S2).

### 3.3 False-failure rate on the correctly constructed set

This set was defined after the failures of Section 3.2 had been attributed, so its rate is an in-sample, post hoc figure, and that few failures fall inside it follows in part from its definition; the cut-off sensitivity below shows how much it depends on that choice. Restricting to X-ray diffraction, resolution ≤ 2.0 Å, no protein in the entry and no deposited covalent bond, with RNA–DNA entries admitted, gave 164 instances, of which 149, from 141 entries and 83 equivalence classes, carried the receptor-clash metric (Table 2). The other 15 have no polymer atom within the 6 Å search radius the check uses, so no contact is measurable; all 15 pass the check. Counting them as passes gives 3 of 164 (1.83%); the reported 2.01% excludes them and is the higher of the two. The rate varied with the resolution cut-off and with which entries were kept (S1 File, Table S8). With the other rules fixed it was 0.00% (0 of 25) at 1.5 Å, 1.14% (1 of 88) at 1.8 Å, 2.01% (3 of 149) at 2.0 Å, 1.44% (3 of 208) at 2.2 Å, 3.37% (10 of 297) at 2.5 Å and 5.22% (29 of 556) at 3.0 Å. Keeping covalently bonded ligands moved the rate more than any cut-off did (8.72%, 15 of 172, at 2.0 Å), and keeping entries that contain protein gave 4.21% (9 of 214).

**Table 2.** False-failure rate of the receptor-clash check on the correctly constructed set.

|  | value |
|---|---|
| false-failure rate at the default threshold of 0.75 | **3 of 149** = **2.01%**, 95% CI 0.00–4.58% |
| PoseBusters protein reference | 1 of 308 = 0.32%, 95% CI 0.06–1.82% |

The three failures were 6CB3/GTP, 4NFO/SPM and 9KL1/SPM, with closest contacts of 2.12, 2.32 and 2.42 Å and one clashing atom pair each. Two of them, 4NFO/SPM and 9KL1/SPM, are spermine, a flexible polyamine. If alternate locations are not matched (Section 2.3), 6C8N/EQ1 also fails, with a closest contact of 0.33 Å across 110 clashing atom pairs, all with atoms of the other alternate location; scored against its own alternate location, its closest contact is 2.99 Å. With mixed alternate locations the clean-set rate would be 4 of 149 (Fisher’s exact P = 0.041 against the protein reference) and the all-but-one threshold 0.684, so the comparison of rates depends on matching alternate locations, which follows their crystallographic meaning.

Failures on deposited poses are treated here as false failures, on the assumption that a deposited high-resolution crystal pose is physically acceptable; a deposited pose can nonetheless carry a genuine modelling error, so this is an upper bound on the check’s error rate. At this sample size the clean-set rate could not be distinguished from the protein reference, which does not establish that the two are equal. Its cluster CI of 0.00–4.58% contains 0.32%, and Fisher’s exact test gave P = 0.104 for 3 of 149 against 1 of 308. No instance was removed from the clean set.

### 3.4 Clash thresholds derived from RNA crystal structures

Three calibration rules were applied to all 149 instances of the clean set (Table 3, Fig 2).

**Fig 2.**
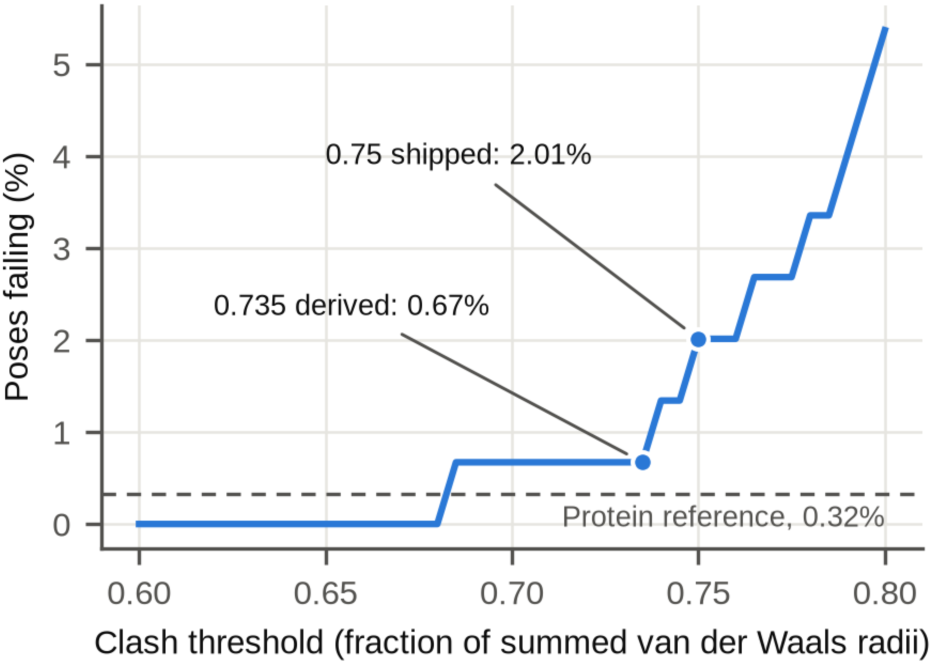
Receptor-clash failure rate as a function of the clash threshold. Correctly constructed set: X-ray, resolution ≤ 2.0 Å, no protein in the entry, no deposited covalent bond (n = 149; each ligand scored against its own alternate location). The threshold is a fraction of the summed van der Waals radii, swept in steps of 0.005. Filled circles, the derived (0.735) and shipped (0.75) thresholds; dashed line, the 0.32% rate (1 of 308) of the PoseBusters protein benchmark.

**Table 3.** Clash thresholds derived from the 149 instances of the clean set.

| **rule** | **threshold** | **failures of 149** |
|---|---|---|
| all crystal ligands but one pass, as applied by PoseBusters | **0.735** (95% CI 0.684–0.788) | 1 |
| protein-matched (1/308 quantile; 149 values cannot give that rate exactly) | 0.709 | 1 |
| 95% rule (0.05 quantile) | 0.798 | 8 |
| shipped threshold | 0.75 | 3 |

All three lie within 0.05 of the shipped threshold of 0.75. The cluster-bootstrap interval of the all-but-one threshold, 0.684–0.788, contains the shipped 0.75. Its lower bound is the smallest relative distance in the set (6CB3/GTP), which becomes the second-smallest value whenever its single-member class is drawn at least twice. The all-but-one threshold is an extreme order statistic, so this interval describes how far the rule moves under resampling rather than the precision of an estimate; the three rules are reported together for that reason.

The all-but-one rule is set by the two most extreme contacts of a set: on the 556 X-ray, no-protein, non-covalent instances at ≤ 3.0 Å it falls to 0.413 (S1 File, Section S17, which also records how PoseBusters took its strain-energy ratio from earlier work [29]).

### 3.5 False-failure rate by resolution and by entry composition

On the X-ray, RNA-only set with covalently bonded ligands removed, the observed failure rate rose as resolution became poorer, from 0.00% below 1.5 Å through 1.83%, 5.30% and 6.74% to 14.77% above 3.0 Å (Fig 3, which gives each bin’s failures and denominator). On the X-ray, no-protein, non-covalent set with RNA–DNA entries admitted and a measurable contact (646 instances, 42 failures, 231 equivalence classes) the rate was 1.14%, 3.28%, 4.73%, 7.34% and 14.44% across the bands up to 1.8, 1.8–2.0, 2.0–2.5, 2.5–3.0 and above 3.0 Å. A logistic trend gave an odds ratio of 3.39 per ångström of poorer resolution (cluster-robust P = 1.4 × 10^−5^), and the rank correlation of band with failure was supported by a permutation of whole classes (ρ = 0.147, P = 2.0 × 10^−4^).

**Fig 3.**
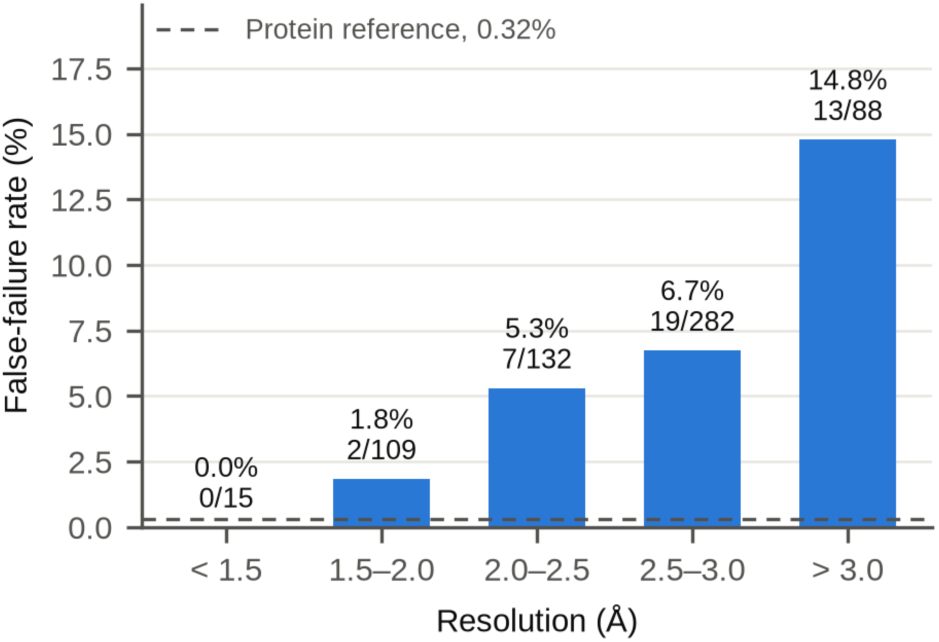
False-failure rate of the receptor-clash check by resolution. X-ray instances from RNA-only entries (no protein or DNA chain), deposited covalent bonds excluded (n = 626). Bins are closed on the left; the 2.5–3.0 Å bin also holds the instances deposited at 3.000–3.009 Å. Bars are labeled with the rate and failures over instances; dashed line, the 0.32% rate (1 of 308) of the PoseBusters protein benchmark.

Among X-ray entries containing protein, non-covalent ligands failed at 16.07% (131 of 815). However, 93 of those 131 failures occurred at resolutions of 3.0 Å or worse (a wider boundary than the above-3.0 Å category of Table 1), and at matched resolution (2.5–3.0 Å) entries containing protein failed at 10.10% against 6.74% for RNA-only entries. Paromomycin, which accounted for 23 of the 131 failures, failed only in structures resolved at 2.9 Å or worse (S1 File, Section S3).

### 3.6 Redocking success with and without the validity requirement

AutoDock Vina 1.2.7 was used to redock the deposited ligand of 139 targets, the X-ray, RNA-only, non-covalent subset of the 173-structure benchmark [7], spanning 71 equivalence classes. All 139 targets docked successfully, and the resulting poses were scored by unmodified PoseBusters (Table 4, Fig 4).

**Fig 4.**
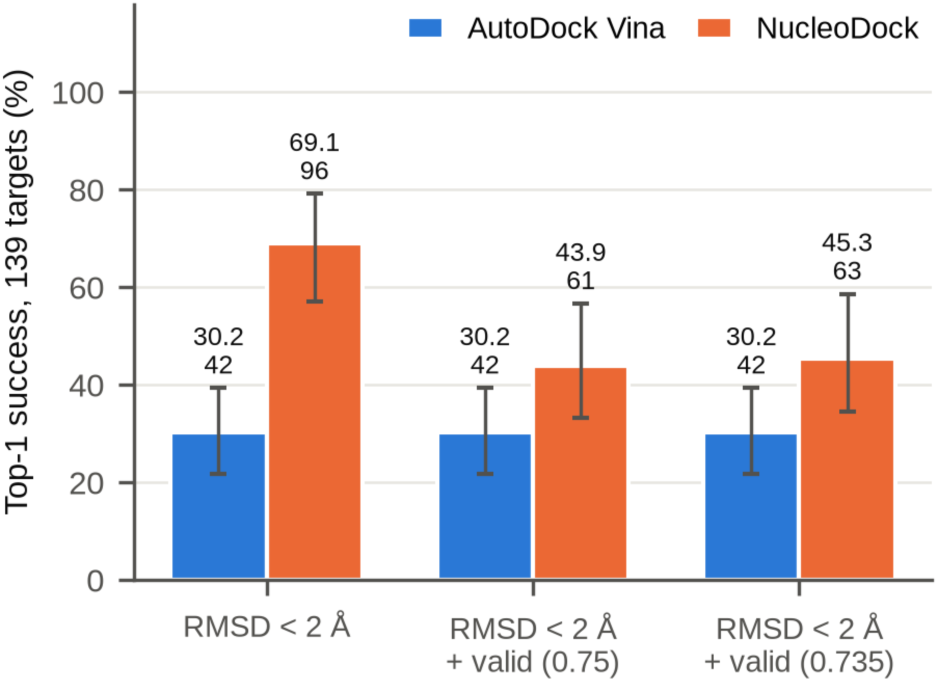
Top-1 redocking success of AutoDock Vina and NucleoDock on the same 139 targets. The X-ray, RNA-only, non-covalent subset of the Agarwal et al. benchmark, redocked into a box defined by the deposited ligand. Success is RMSD < 2.0 Å, measured in the crystal frame without superposition, alone or with PoseBusters validity at the shipped (0.75) or derived (0.735) threshold. Bars are labeled with the rate (%) and the number of successes; whiskers, 95% confidence intervals from a bootstrap over 71 equivalence classes (4,000 resamples). RMSD, root-mean-square deviation.

Accuracy was measured in the crystal frame, without superposition (Section 2.4). On that measure 42 of the 139 AutoDock Vina poses (30.22%; 95% CI 21.74–39.29) lay within 2.0 Å of the deposited ligand, against 95 (68.35%) after optimal superposition: 53 poses had the shape of the deposited ligand but lay elsewhere in the search box (median RMSD 4.43 Å unaligned, 1.27 Å aligned). Rates of success in this paper use the unaligned value only. The AutoDock Vina poses placed 95 ligands within 2 Å after superposition; an earlier run with the same settings gave 96, but its poses were not archived.

Requiring physical validity cost AutoDock Vina no accurate pose: all 42 were valid, so the two thresholds gave the same rate (30.22%). The minimum relative distance to the polymer across all 139 AutoDock Vina poses was 0.753, so no AutoDock Vina pose failed the receptor-clash check at either threshold.

Of the 139 poses, 132 (95.0%) were PoseBusters-valid. Seven poses were invalid, none accurate (all lay 4.4 Å or more from the deposited ligand): six failed the internal-energy check (1EVV/SPM, 1FUF/SPM, 1XPF/SPM, 2O3Y/ SPM, 3GX3/SAH and 4P5J/SPM) and one the flatness of a non-aromatic ring (2BE0/JS5).

This experiment is redocking into the ligand-defined box, the regime in which Agarwal et al. [7] report AutoDock Vina near 63%. The superposed value here (68.35%) lies in that region and the unaligned value (30.22%) does not; which RMSD convention that benchmark used was not checked here, so the two should not be read as a replication. Neither is comparable to the approximately 27% that the same paper reports for a large blind box.

### 3.7 Validity of the deposited ligands of the docking set

The deposited ligand of each of the 139 targets was scored by the same PoseBusters configuration. This gives a reference validity rate for the deposited poses, not an upper bound on docking success. The deposited ligand failed at least one check on 10 of 139 targets (7.19%), so 92.81% of targets have a deposited ligand that passes every check. This is a reference rate, not a strict ceiling, because a docked pose can pass where the deposited pose fails. The receptor-clash check failed 7 targets at the shipped 0.75 (5.04%; 95% cluster CI 0.96–9.70) and 4 at 0.735 (2.88%; 0.00–7.64). The other failing checks were the distance to organic cofactors (3 targets), the volume overlap with organic cofactors (2) and the flatness of one non-aromatic ring (1); a target can fail more than one check, and no chemistry check failed. The validity-qualified success of both programs lies below 92.81% (AutoDock Vina 30.22%, NucleoDock 43.88%), so failures of the deposited ligands do not explain the gap; NucleoDock poses failed the receptor-clash check on 53 targets, against 7 for the deposited ligands.

### 3.8 Paired comparison with a nucleic-acid-specific docking program

NucleoDock [18], the nucleic-acid-specific program whose exclusion of checks motivated this study, was run from its released code on the same 139 targets, into the same deposited-ligand-defined box, and its top-1 pose was scored by the same unmodified PoseBusters configuration (Tables 4 and 5, Fig 4). Both programs returned a pose for every target, so the comparison is paired, target for target.

**Table 4.** Top-1 redocking success of AutoDock Vina and NucleoDock over the same 139 targets. Rates with the number of successes and the 95% CI from a bootstrap over the 71 equivalence classes.

| **criterion** | **AutoDock Vina, % (n of 139)** | **95% CI** | **NucleoDock, % (n of 139)** | **95% CI** |
|---|---|---|---|---|
| RMSD < 2.0 Å only | 30.22 (42) | 21.74–39.29 | 69.06 (96) | 57.04–79.09 |
| RMSD < 2.0 Å, valid at 0.75 (shipped) | 30.22 (42) | 21.74–39.29 | 43.88 (61) | 33.13–56.57 |
| RMSD < 2.0 Å, valid at 0.735 (derived) | 30.22 (42) | 21.74–39.29 | 45.32 (63) | 34.56–58.57 |

**Table 5.** Paired difference in top-1 redocking success, NucleoDock minus AutoDock Vina. Cluster intervals and permutation tests resample the 71 equivalence classes; McNemar p values treat the 139 targets as independent and are shown for reference. pp, percentage points.

| **criterion** | **difference, pp (95% cluster CI)** | **cluster permutation p** | **discordant targets (Vina only / NucleoDock only)** | **McNemar p** |
|---|---|---|---|---|
| RMSD < 2.0 Å only | **38.85** (26.47 to 50.00) | < 0.001 | 64 (5 / 59) | 9.0 × 10<sup>−13</sup> |
| RMSD < 2.0 Å, valid at 0.75 (shipped) | 13.67 (1.34 to 27.52) | 0.053 | 49 (15 / 34) | 0.0094 |
| RMSD < 2.0 Å, valid at 0.735 (derived) | 15.11 (2.42 to 29.20) | 0.030 | 51 (15 / 36) | 0.0046 |

By RMSD alone, NucleoDock placed 54 more ligands within 2 Å than AutoDock Vina, 69.06% against 30.22%. The lead was 38.85 percentage points (95% CI 26.47–50.00, bootstrapped over equivalence classes; cluster permutation p < 0.001). Requiring the same poses to be physically valid cost NucleoDock 25.18 points (95% CI 14.04–35.33) and AutoDock Vina none (all 42 accurate poses were valid), so the difference between the programs shifted by 25.18 points (95% CI 14.04–35.33) toward AutoDock Vina. With validity required, NucleoDock still led, by 13.67 points at 0.75 (95% CI 1.34–27.52; cluster permutation p = 0.053) and by 15.11 at 0.735 (2.42–29.20; p = 0.030). McNemar’s exact test, which treats targets as independent, gave p = 0.0094 and 0.0046. With validity required, the lead fell from 38.85 to 13.67 points (95% CI 1.34 to 27.52); the cluster permutation test gave P = 0.053, so the validity-qualified lead is not distinguishable from zero at the 0.05 level.

On the 37 targets that both programs placed within 2.0 Å, all 37 AutoDock Vina poses were valid, against 27 NucleoDock poses at 0.75 and at 0.735.

Of the 96 accurate NucleoDock poses, 35 were not valid at 0.75. The receptor-clash check accounted for 28 of them, followed by internal energy (6), the organic-cofactor distance (1), the inorganic-cofactor distance (1) and the internal steric clash (1); one pose may fail more than one check, and the volume overlap with the polymer failed none of the accurate poses. The receptor-clash check therefore failed NucleoDock poses, whereas it failed no AutoDock Vina pose (Section 3.6), and the two thresholds no longer gave identical rates: the minimum relative distance to the polymer was 0.199 over the NucleoDock poses against 0.753 over the AutoDock Vina poses, and two accurate NucleoDock poses lay between 0.735 and 0.75.

Ligand geometry may contribute to the difference, because AutoDock Vina started from the deposited ligand and NucleoDock built the ligand from SMILES (Section 2.4). Ligand geometry acts most directly on the intramolecular checks, which 7 of the 35 accurate-but-invalid NucleoDock poses fail (internal energy 6, internal steric clash 1). Its effect on the receptor-clash failures, 28 of them in accurate poses, is examined in Section 3.9.

Counted over all 139 top-1 poses, 63 NucleoDock poses (35 of them accurate) failed at least one check, against 7 AutoDock Vina poses (none accurate). The receptor-clash check at 0.75 failed 53 NucleoDock poses (28 accurate), and 33 of those (26 accurate) failed no other check. The internal-energy ratio failed 13 NucleoDock poses (6 accurate), 4 of them with no other failed check, and the volume overlap with the receptor failed 13 (none accurate), each together with another check. AutoDock Vina lost poses to the internal-energy ratio (6, none accurate) and one ring flatness, and none to the receptor clash. The receptor-clash check therefore accounts for most of the validity cost of NucleoDock, although other checks also exclude some of its poses.

### 3.9 The receptor-clash failures of NucleoDock

The receptor-clash failures follow the NucleoDock pose and not the target. None of the 53 NucleoDock clash failures also failed the check for the AutoDock Vina pose of the same target, and 5 also failed it for the deposited ligand. The most severe pose (4YAZ, ligand 4BW) places a ligand carbon 0.655 Å from a receptor nitrogen (relative distance 0.199); the median failing pose had 2 clashing atom pairs. The ratio of a pose’s internal energy to that of a relaxed conformer ensemble had a median of 2.88 for AutoDock Vina, 2.30 for NucleoDock and 1.72 for the deposited ligands. These medians do not show that SMILES-derived geometry contributes to the receptor clashes (S1 File, Section S15). A matched-input comparison follows in Section 3.10.

### 3.10 Robustness of the comparison to ligand input, program, repeat and training overlap

Each analysis below retains the direction of the main result: NucleoDock leads on RMSD and, with validity required, keeps a smaller lead whose size depends on the comparison; the full text, intervals and tables are in S1 File, Sections S12 to S14 and S16.

#### Matched ligand input

AutoDock Vina docked from an RDKit conformer of the SMILES string that NucleoDock received (Section 2.7) placed 32 of 139 ligands within 2.0 Å (23.02%), all of them valid, with no receptor-clash failure (minimum relative distance 0.809). NucleoDock led by 46.04 points on RMSD (95% cluster CI 30.95–58.00) and by 20.86 points with validity required at 0.75 (10.42–30.91; P = 0.001). Starting from the RDKit conformer instead of the deposited ligand lowered the AutoDock Vina rate by 7.19 points (−15.76 to 1.64), so the unequal ligand input does not account for the lead, although the interval does not exclude a drop of up to 15.76 points.

#### A third program

smina 2017.11.9 docked the deposited ligands. Five macrocyclic ligands that it cannot read were counted as failures to dock; on that basis it placed 40 ligands within 2.0 Å (28.78%), all valid, with no receptor-clash failure. NucleoDock led by 40.29 points on RMSD (28.93–50.72) and by 15.11 with validity required (3.08–28.57; P = 0.023). With the five prepared as rigid rings the figures were 42 poses (30.22%), a lead of 38.85 points (26.36–50.00) and, with validity required, 13.67 points (1.34–27.35; P = 0.049). Requiring validity cost none of the AutoDock Vina and smina arms an accurate pose, against 25.18 points for NucleoDock.

#### Repeat of the NucleoDock run

This repeat, with the seed fixed by the program, tests the reproducibility of the execution and not the variability across random seeds. Reproducibility of the NucleoDock run was assessed on the superposed RMSD, which was computed separately from the poses of each run; the unaligned RMSD of the first run is read from the poses of the repeat, except for the three targets nearest the 2 Å cut. Repeating the 139 runs with the same container digest, configuration and inputs gave 125 poses within 2.0 Å after superposition against 126, identical to 0.001 Å on 119 of 139 targets, with a largest change of 0.48 Å. Without superposition the repeat placed 96 ligands within 2.0 Å (69.06%) and 62 with a valid pose (44.60%), against 61 in the first run: one validity-qualified call changed, a gain, and the validity check alone changed on 3 targets (1 gained, 2 lost). The receptor-clash check failed 55 poses against 53 and the validity cost was 24.46 points against 25.18. The runs differed by 0.72 points on the validity-qualified criterion, small against the 13.67-point lead and the 25.18-point cost.

#### Overlap with the development data of NucleoDock

The public repository lists no training, validation or test sets. The preprint’s 125-complex external test set, which it excludes from training, contains 53 of the 139 targets; the use of the other 86 in training can be neither confirmed nor excluded. On the 53 targets NucleoDock led AutoDock Vina by 39.62 points on RMSD (20.93–55.56) and by 5.66 with validity required (−15.79 to 27.69); on the other 86 the differences were 38.37 (24.36–51.69) and 18.60 (2.33–36.36). Its RMSD success was 67.92% on the first subset and 69.77% on the second, which gives no sign of a contribution from training overlap, although the test cannot exclude one. The lead on RMSD is present in both subsets; with validity required it is present on the 86 and not distinguishable from zero on the 53, where the interval is wide.

## 4. Discussion

On a high-resolution, non-covalent, protein-free set of 149 poses, the protein-calibrated receptor-clash threshold failed 3 poses (2.01%), a rate these data cannot distinguish from the protein reference, and failures under naive application were concentrated in covalently bonded nucleotides, entries that contain protein, cryo-EM entries and poorly resolved structures. The protocol and the clash threshold support it. The docking comparison (Section 4.6) shows how large the validity requirement can be for one program; it rests on two programs and a borderline test, and is supporting evidence for reporting validity.

### 4.1 Sources of failure under naive application

On the correctly constructed set the receptor-clash check failed 3 of 149 deposited poses, a rate that could not be distinguished from the protein reference. The large failure rate under naive application coincides with properties of the input set: covalently bonded nucleotides, experimental method and resolution, which overlap with entries containing protein.

The two rates, 18.0% and 2.01%, come from sets that differ in composition. Nucleotides covalently bonded into the chain but deposited as free ligands contributed 23.0% of failures, and all but 7 of the rest came from entries containing protein, cryo-electron microscopy or X-ray structures worse than 2.0 Å, classes that largely coincide (Section 3.2). In the covalent class the check is not failing; it is reporting a bonded contact as a clash, which is the correct response once the nucleotide is presented as a free ligand. The 18.0% is therefore a rate for naive preparation of deposited structures, a catalogue of construction pitfalls, and not an RNA-specific failure rate of the check. The remaining failure rate rose as resolution became poorer, so a headline failure rate for RNA depends on how much low-resolution data the set admits (Section 3.5). Paromomycin shows this within one compound (Section 3.5). Alternate locations are a smaller hazard of the same kind: mixing them produces three receptor-clash failures, one of them the most severe in the clean set, and nothing in the resulting receptor file shows the error.

### 4.2 Clash thresholds

The thresholds derived here do not establish whether an RNA-specific calibration is needed, and on these 149 instances they give no indication that one is. All three rules returned values within 0.05 of the shipped 0.75, and the rule PoseBusters itself used returned 0.735, with a cluster-bootstrap 95% CI of 0.684–0.788 that contains 0.75. The false-failure rate at the shipped threshold, 2.01%, could not be distinguished from the protein benchmark’s 0.32% (cluster CI 0.00–4.58%; Fisher’s exact P = 0.104), with no instance excluded. The counts are small: the interval extends to about 4.6%, so a small RNA-specific excess cannot be excluded. The derived thresholds and the observed rates are compatible with the shipped threshold on data of this kind, and neither excludes a small RNA-specific excess. The all-but-one threshold was derived and evaluated on the same 149 instances, so 0.735 is a descriptive in-sample value and its out-of-sample calibration is untested. PoseBusters applied that all-but-one rule, which two extreme contacts can move (Section 3.4), to 308 hand-curated reference complexes. The RNA set here was not curated by hand, so all three rules are reported, and the threshold is given with its interval.

### 4.3 Assignment of receptor atoms and exclusion of checks for nucleic acids

PoseBusters assigns receptor atoms to checks by record type, so an RNA chain written as ATOM records is scored by the receptor-clash check exactly as a protein chain is, and modified nucleotides written as HETATM records are scored by the organic-cofactor check (Section 2.4). In the clean set this changed no result, but analyses that report only `minimum_distance_to_protein` omit those contacts: 693 of the 2,970 instances had a modified nucleotide in the pocket and 451 had one within 4.0 Å of the ligand (S1 File, Section S4). Comparisons with published PoseBusters rates should state the version used, because results produced before the fix of issue #96 counted organic cofactors in the receptor-clash check.

These results bear on whether checks need to be excluded for nucleic acids. No check was disabled when scoring the deposited poses, and the check-by-check pass rates below show where the losses fall; the threshold analysis, however, covers the receptor-clash check alone, so the evidence is strongest for that check. On RNA the validity loss is mostly intermolecular: on the X-ray, RNA-only, non-covalent set at ≤ 3.0 Å, the chemistry checks passed at 99.5%, the intramolecular checks at 98.4% and the intermolecular checks (receptor and cofactor distances, volume overlaps and the maximum ligand–receptor distance) at 89.9%. On the AutoDock Vina output the receptor-clash check failed no pose, and keeping every check cost no accurate pose (0.00 percentage points). On the NucleoDock output it failed many: the check that study advises interpreting conservatively accounted for 28 of its 35 accurate-but-invalid poses, and keeping every check cost 25.18 points (the repeat gave 24.46 points; Section 3.10). The same check passed all but 3 of the 149 deposited crystal poses in the clean set, although that comparison alone does not establish what produced the failures among the docked poses. The radical species given as an example for excluding checks were not observed on the chemistry supplied by the RCSB Protein Data Bank: the radical check passed on all 2,969 retained ligands, including 128 instances from 119 of the 125 entries of the NucleoDock set. Components that HARIBOSS did not list and the additive filter of Section 2.2 excluded, such as ions, were not tested. These results concern deposited structures and redocked poses prepared as described here. They do not show what produced the failures in that study, whose excluded checks are not stated.

The two kinds of evidence support different statements. The deposited crystal poses show that the receptor-clash check at its shipped threshold rarely fails a well-constructed RNA complex (2.01%, interval to 4.58%), so on these data the check rarely fails a well-constructed deposited complex. The docking arms show that the poses of one program fail it often and those of two general-purpose programs do not, which points to the poses and not to the calibration. Neither result shows that the exclusion of checks for nucleic acids was or was not justified in the study that proposed it.

### 4.4 Protocol for applying PoseBusters to nucleic-acid complexes

1. Resolve each ligand by instance: chemical component identifier, chain **and** residue number. Without the residue number, a query can return a different copy of the ligand; for 6U7Y/C5P the copy returned lay 34.8 Å away.
2. Exclude ligands carrying a deposited `struct_conn covale` bond to the polymer, using the deposited annotation rather than a distance cutoff. Excluding covalent complexes follows existing practice for nucleic-acid docking sets [8].
3. Strip the ligand from the receptor by entity and instance, never by residue name. A chemical component identifier can equal a polymer residue name (`N`, `C`, `G`); name-based stripping removes polymer atoms, up to 80 per case here and, for 8P2K, the entire receptor.
4. Keep only receptor atoms with a blank alternate location or the ligand’s own. A ligand in alternate location B scored against atoms in A produces a clash of 110 atom pairs at 0.33 Å on a 1.9 Å structure (6C8N/ EQ1); 35 of 2,970 receptors mixed alternate locations this way.
5. Report entries containing protein separately from RNA-only entries.
6. Stratify by resolution and experimental method, and do not pool X-ray diffraction with electron microscopy.
7. Run PoseBusters unmodified, at its default thresholds, and report the results as such. About 2% false failures were observed on high-resolution data (3 of 149 here), for the receptor-clash check, which is the only check this study calibrates.

### 4.5 Implementation

Steps 1 to 4 of the protocol fail silently when done incorrectly, so they are implemented in `rnaprep` 0.3.0. It takes a PDB entry and one ligand instance, refuses a covalently bonded ligand, an empty pocket and a receptor with no polymer atoms, and writes the receptor PDB file and ligand SDF file that PoseBusters reads. The mapping of protocol steps to operations is given in S1 File, Section S5.

### 4.6 Validity and the comparison of docking programs

On the same 139 targets, a nucleic-acid-specific program led a general-purpose one by 38.85 percentage points on RMSD alone, measured without superposition, and requiring physical validity shrank that lead by 25.18 points (95% CI 14.04–35.33) to 13.67 (95% CI 1.34–27.52; Section 3.8). The lead was not reversed; whether any lead remains is not settled by these data (cluster permutation P = 0.053, and the interval includes zero under another clustering; Section 3.10). The validity cost was program-dependent, at none for AutoDock Vina, whose 42 accurate poses were all valid, and 25.18 points for NucleoDock, so, for these two programs, the RMSD-only lead of 38.85 points was about three times the validity-qualified lead of 13.67 points. RNA docking benchmarks should therefore report that rate alongside the RMSD-only rate. The evidence is narrower than the audit of Sections 3.1 to 3.5: it is two programs, one redocking regime, a validity-qualified lead whose permutation test is borderline (P = 0.053) and whose interval includes zero when the 24 unclassed targets are one cluster (Section 3.10), and a validity cost that may reflect the absence of a physics-based refinement step in the NucleoDock output, which AutoDock Vina and smina perform against a steric repulsion term, and not the quality of the underlying model. It shows that the cost can be large and program-dependent, which is the reason to report it, and it is not a ranking. The pattern held when AutoDock Vina started from the ligand geometry that NucleoDock received and against smina, a third program, and on the targets inside and outside NucleoDock’s external test set (Section 3.10). Superposition before scoring, which measures shape and not placement, classified 53 AutoDock Vina poses and 29 NucleoDock poses as accurate that lie in the wrong place in the receptor frame, so accuracy here is stated in the crystal frame. Benchmarks of protein–ligand docking and co-folding programs likewise find that success falls on targets unlike the training data [30,31], which is why the overlap of the targets with the training data of a learned program has to be stated.

### 4.7 Limitations

- Accuracy is the RMSD of the top-1 pose to the deposited ligand in the receptor frame, without superposition, and a pose within 2 Å is counted as accurate whatever the receptor contacts. Two provenance limits apply. The AutoDock Vina poses were regenerated with the protocol of the first run because the poses of the first run were not archived (95 against 96 poses within 2 Å after superposition), and the PoseBusters report was recomputed for them. The first NucleoDock run’s poses were not archived either; its unaligned RMSD is read from the poses of its repeat, which reproduce its superposed RMSD to 0.001 Å on 119 of 139 targets, and from the first run’s own poses for the three targets whose RMSD lies nearest the 2 Å cut (2GDI, 2HOJ and 2HOM), while the other 17 targets whose superposed RMSD changed between the runs lie at least 0.5 Å from the cut in the repeat’s unaligned RMSD, and no superposed RMSD changed by more than 0.49 Å. The NucleoDock RMSD success here (69.06%) is above the 56.0% top-1 success that its preprint reports on its own 125-complex test set; the box defined by the deposited ligand, the possible training overlap and the CPU execution may each contribute, and the comparison does not separate them.
- The clean set was defined after the failures had been attributed to input categories (Section 3.2), so its 2.01% is an in-sample figure, and it moves with the choices (Section 3.3, cut-off curve). It is small, at 149 poses over 83 equivalence classes, and its confidence interval of 0.00–4.58% is wide. Only the receptor-clash check was recalibrated; the other checks keep their protein thresholds. A larger high-resolution RNA–ligand set is needed to establish whether the false-failure rate on RNA equals that on protein or is a few percentage points higher.
- Solution-NMR entries were evaluated on model 1 only; their failure rate without protein or covalent ligands, 4.82%, is reported but not interpreted. Ligand selection blends the curation of HARIBOSS with an additive filter that admitted some additives (6DTD/CIT is citrate).
- The docking comparison is scoped to NucleoDock against AutoDock Vina, with smina as a second general-purpose program (Section 3.10), and to redocking into a box defined by the deposited ligand, the regime of the source benchmark and the easiest docking task. No program was tuned, and the result is not a general ranking. RNA changes conformation on ligand binding [32], so a cross-docking or blind-box comparison could show a different validity cost and is the next experiment. Co-folding programs [13–16] were not run.
- The two programs received the ligand differently (AutoDock Vina the deposited structure, NucleoDock SMILES), which may favor AutoDock Vina on the intramolecular checks, failed by 7 of the 35 accurate-but-invalid NucleoDock poses. The receptor-clash cost is not explained by this difference (Sections 3.9 and 3.10), but NucleoDock started from the deposited conformer was not run, so the matched-input test is one-sided.
- NucleoDock exposes no seed: the shipped configuration fixes it at 42 and the pose-generation stage ran on the CPU. A repeat of all 139 targets changed 1 superposed RMSD call and 1 validity-qualified call at 0.75 (Section 3.10), so the 25.18-point validity cost rests on a run that a repeat reproduces to within about one target; the repeat also differed from the first run in scheduling and nodes. The AutoDock Vina and smina arms used one search seed per target. The cost comes from one commit of the released code, and the rates NucleoDock reports were produced with an unstated subset of PoseBusters checks, so ours are not directly comparable to them.
- Whether NucleoDock was trained on the 86 targets outside its external test set cannot be determined from the public repository. Programs that learn from deposited complexes reproduce training poses [31], so an overlap would raise their RMSD success on the overlapping targets; the success rate was the same inside and outside the test set (Section 3.10).
- NucleoDock was scored on the first of the ten poses it writes per ligand, matching the top-1 rates it reports; a top-N or ensemble criterion would give different rates for both programs.
- The alternate-location rule of Section 2.3 only removes receptor atoms, so it can only turn clash-check failures into passes; passing instances were not audited for construction errors that would hide a clash. Of the 2,969 instances checked, 38 had no polymer atom within the range of the receptor-clash check and are excluded from receptor-clash rates, and 8P2K/N could not be checked at all (Section 2.3).

## 5. Conclusions

Applied to deposited RNA–ligand structures without filtering the input, PoseBusters’ receptor-clash check failed 18.0% of poses, and most of those failures came from covalently bonded nucleotides presented as free ligands, entries that contain protein, cryo-electron microscopy and poor resolution. On 149 high-resolution crystal poses from RNA and RNA–DNA entries, built with the protocol given here and selected after the failures were attributed, the failure rate was 2.01% and could not be distinguished from the protein reference, although the interval does not exclude a modest excess; the rule PoseBusters used selected 0.735 against the shipped 0.75. No check was excluded. We provide a construction protocol and an implementation of its first four steps; no new threshold set is proposed. In a paired comparison on 139 redocking targets, with accuracy measured without superposition, one program led another by 38.9 points on RMSD alone and requiring validity cut that lead to 13.7 points, a lead not clearly distinguishable from zero (P = 0.053), because the leading program lost 25.2 points to the check and the other lost none. The lead remained, with a smaller size, when AutoDock Vina started from the ligand geometry that NucleoDock received, against smina, and on the targets outside NucleoDock’s external test set; with validity required its interval excludes zero in most of these comparisons and includes it on the 53 targets inside the test set. For these two programs, in one redocking regime with one run per program, the comparison supports reporting physical validity alongside RMSD in RNA docking benchmarks on inputs prepared as the protocol describes; it does not establish a ranking or show how general the validity cost is.

## Data availability

The code, data tables, preparation tool and container recipes are archived on Zenodo, https://doi.org/10.5281/zenodo.24320791 (concept DOI, which resolves to the latest version of the archive), under CC BY 4.0, and can be downloaded without an account. No private service is needed to reproduce the results: PoseBusters 0.6.5 and every Python dependency are pinned in a hash-locked requirements file, the `rnaprep` container can be rebuilt from the archived Dockerfile, and NucleoDock is public at https://github.com/ ShiYue3384/NucleoDock. The archive contains the per-instance results with every check and raw metric for the 2,969 checked instances, the input set, the per-check false-failure rates by stratum, the redocking results of both programs with the paired comparison, every pipeline step from the PDB query to the figures, and `rnaprep` 0.3.0 with its Dockerfile and Nextflow wrapper; the file-by-file list is in S1 File, Section S6. Accuracy is rescored from the committed pose archives (`review/gap-closure-2026-11-20/ new_arms/poses_*.tar.gz`) and the deposited ligand coordinates of the prepared inputs (`inputs_prepared_pdbqt.tar.gz`) by `rescore_rmsd_arms.py` and `score_arm_from_pdbqt.py`, so no external download is needed. `headline_numbers.py` and `methods_numbers.py` recompute the quoted numbers listed in Statistical analysis and fail on any difference, and `make_figures.py` regenerates Figs 1–4. These scripts run offline in an environment pinned by a hash-locked requirements file for Python 3.12; the upstream steps, which need network access, have a separate pinned requirements file.

The 139 NucleoDock runs and their repeat, and a digest-pinned PoseBusters reproduction of the first AutoDock Vina run (133 of 139 poses passed all 22 checks, none discordant from the local run; the regenerated poses of the version of record gave 132 of 139), were executed on the Omic platform; the AutoDock Vina arms, the matched-input arm and the smina arm were run on a local workstation. `rnaprep` 0.3.0 has been run locally only, and no platform run of it covers the full set of 2,970 instances. Run identifiers, commits, container digests and the platform and local run details are in S1 File, Section S6.

Structures are available from the RCSB PDB under the entry identifiers given; PDB data are released under the CC0 1.0 Public Domain Dedication. The HARIBOSS export used to select ligands is distributed by HARIBOSS under CC0 1.0 and is redistributed with attribution [19]. The two benchmark sets are redistributed as lists of PDB identifiers taken from the supporting information of the source papers [7,18].

## Supporting information

**S1 File. Supporting information.** Section S1, alternate-location handling; Section S2, attribution of receptor-clash failures under a second order of causes; Section S3, receptor-clash failures of paromomycin by resolution; Section S4, polymer residues written as HETATM records and the radical check; Section S5, the `rnaprep` operations; Section S6, archive contents and run details; Section S7, the literature and citation screen; Sections S8 to S14, the clean-set sensitivity table, the regression, the checks that fail the docked poses and the deposited ligands, and the results of the additional docking arms; Section S15, the receptor-clash failures of NucleoDock; Section S16, the robustness analyses in full; Section S17, the all-but-one rule and the PoseBusters calibration.

## Acknowledgments

The research was carried out on the Omic research platform (https:// omic.ai), which ran the docking, validity-checking and structure-preparation workflows, stored their inputs and outputs, and recorded every run.

