## Supplementary material for "Input Construction Accounts for Most PoseBusters Clash Failures on RNA–Ligand Complexes": S1 File. Supporting information

Section numbers in the main text refer to the manuscript. Every number below is produced by the scripts named in the Data availability section of the manuscript.

#### S1. Alternate-location handling

Each receptor kept only atoms with a blank alternate location or the ligand's own. A shared label does not by itself show that conformers in different residues are correlated, so this rule assumes the deposited labeling is consistent within the pocket; where it is not, the rule removes receptor atoms that could have been retained, which can only turn clash-check failures into passes. In 6C8N, for example, EQ1 is modeled in alternate location B at occupancy 0.40 and the nucleotide G15, which it overlaps, only in alternate location A at 0.60, so the two are not modeled together. For each of the 2,970 instances, the alternate location of the scored ligand atoms was read from the pocket mmCIF file; 56 ligands sat in a non-blank alternate location, and 35 of their receptors also held atoms in another one, which were removed. Matching alternate locations changes the receptor-clash outcome of three instances (6C8N/EQ1, 5SWD/ADE and 6P1U/CTP) and the organic-cofactor outcome of four, all from fail to pass. In none of the 56 instances did the scored ligand atoms span two alternate locations.

#### S2. Attribution of receptor-clash failures under a second order of causes

The causes of Table 1 overlap, so the shares depend on the order in which they are tested. Under an order that tests method and resolution before protein, the same 527 failures split into 121 covalent, 229 electron microscopy, 102 X-ray above 3.0 Å, 62 X-ray at 2.0–3.0 Å, 6 attributable only to protein in the entry and the same 7 residual (121 + 229 + 102 + 62 + 6 + 7 = 527). The covalent share and the 7 residual failures do not depend on the order.

**Further detail on the residual and covalent classes.** The seven residual failures were four solution-NMR entries (1RAW, 1Q8N, 2KGP and 7FJ0), two

X-ray structures at ≤ 2.0 Å with no protein in the entry (6CB3 and 4NFO) and one RNA–DNA hybrid at 1.60 Å (9KL1). The covalently bonded class includes 5′-terminal nucleotides: entries 8KEB and 9LKE carry deposited `covale` records joining G2 P to GTP O3′ at 1.55 and 1.56 Å, which is a phosphodiester bond length. The receptor-clash check flags these contacts because the nucleotide is presented to it as a free ligand. Covalent annotation and interatomic distance did not agree, so neither can substitute for the other: 87 instances had a polymer contact below 1.8 Å with no deposited `covale` record, and 45 instances carried a record without a short contact. Measured in the ligand's own alternate location (Section 2.3), 67 of the 87 instances had a short contact without a record; the other 20 short contacts were with atoms of another alternate location.

### S3. Receptor-clash failures of paromomycin by resolution

Paromomycin accounted for 23 of the 131 receptor-clash failures among X-ray entries containing protein, and its failures were concentrated at poorer resolution. Across all 101 paromomycin instances carrying the metric, the median resolution was 3.14 Å. None of the 14 instances at ≤ 2.5 Å failed, whereas 32.8% of the 67 instances above 3.0 Å failed. All 26 paromomycin failures across the 101 instances, the 23 in entries containing protein among them, were in structures resolved at 2.9 Å or worse.

### S4. Polymer residues written as HETATM records

Of 2,970 instances, 693 had at least one residue that the deposited mmCIF file assigns to a polymer entity and that is written as a HETATM record in the pocket, and 451 had one within 4.0 Å of the ligand. The most frequent were PSU (186 instances), 2MA (143), 5MC (130), UR3 (107), MA6 (81) and OMG (77). In the correctly constructed set, 54 instances had such a residue and 47 had one within 4.0 Å of the ligand; none failed the organic-cofactor clash check under the default thresholds.

**Radical check.** The radical check passed on all 2,969 deposited ligands checked, with bond orders taken from the SDF supplied by the RCSB Protein Data Bank.

### S5. The rnaprep operations

`rnaprep` 0.3.0 takes a PDB entry and one ligand instance and writes the receptor PDB file and ligand SDF file that PoseBusters reads:

```
rnaprep --pdb-id 6CB3 --ligand GTP --chain A --seq 219 --
outdir out/
```

Each protocol step of Section 4.4 maps to one operation:

- Step 1: the ligand is selected by chemical component identifier, `auth_asym_id` and `auth_seq_id`.
- Step 2: the deposited `struct_conn` records are read, and a covalently bonded ligand is refused unless `--allow-covalent` is given.
- Step 3: the ligand is removed by entity and instance, and polymer residues of the same name are kept.
- Step 4: only receptor atoms with a blank or the ligand's own alternate location are kept.

Two guards reject inputs that would otherwise pass unnoticed. The RCSB ModelServer answers a query with a wrong residue number with HTTP 200 and a header-only mmCIF file, so `rnaprep` rejects an empty pocket rather than writing it out. It also rejects a receptor with no polymer atoms, because PoseBusters raises an exception on an empty receptor instead of reporting a failed load. A Dockerfile and a Nextflow wrapper are included.

### S6. Archive contents

The archive contains the per-instance results with every check and raw metric (`per_instance_checks.csv`, 2,969 checked instances), built by `build_per_instance_checks.py` from the full PoseBusters run (`unified_checks.csv`, 2,970 rows), the rescoring of the seven name-collision receptors (`rerun_name_collisions.py`) and of the 35 receptors restricted to the ligand's own alternate location (`rerun_altloc.py`, which also lists them); the input set (`unified_instances.csv`); per-check false-failure rates by stratum (`false_failure_rates.csv`, built by `build_false_failure_rates.py`); the polymer HETATM table; the redocking results of both programs (`a_vina_checks.csv` and `b_nucleodock_checks.csv`, identical columns, built by `run_vina_redock.py` and `score_nucleodock.py` for the first scoring, and in the version of record by `rescore_rmsd_arms.py` and `score_arm_from_pdbqt.py` from the pose archives; `rmsd_top1` is the RMSD in the crystal frame without superposition, `rmsd_aligned_top1` the superposed RMSD kept for discussion, rmsd_aligned_original_run the value the first scoring recorded, and rmsd_pose_source where each row's poses came from) with the paired comparison, its cluster-level inference (paired_cluster_bootstrap.csv, built by paired_cluster_bootstrap.py) and the per-program rate tables (docker_comparison.csv, docker_comparison_per_target.csv, built by compare_dockers.py, and plan_a_results.csv, plan_a_per_target.csv, plan_a_results_nucleodock.csv and plan_a_per_target_nucleodock.csv, built by analyze_plan_a.py, one table per program so that no rate can be read off the wrong run); the NucleoDock input builder, uploader and batch driver (build_nucleodock_inputs.py, upload_nucleodock_inputs.sh, nd_batch.mjs) with the run identifiers of all 139 runs; every pipeline step from the PDB query to the figures; and rnaprep 0.3.0 with its staging script, Dockerfile and Nextflow wrapper.

The archive also carries the analyses that address the reviewers' further points: the validity ceiling of the deposited ligands, the per-check costs, the clash diagnosis, the sensitivity of the clean-set rate to the cut-off, the regression and the resolution trend (review_gaps.py); the matched-input AutoDock Vina arm and the smina arm, with their prepared inputs, poses, scored tables and rate tables (prep_gap_inputs.py, prep_gap_rigid_dep.py, run_gap_dock_local.py, run_gap_smina_docker.sh, score_gap_arms.py, merge_scored.py, gap_arms_all.sh, gap_arms_analysis.py); the overlap analysis with the external test set of NucleoDock; and gap_numbers.py, which recomputes the numbers these analyses add to the text.

**Platform and local runs.** The 139 NucleoDock runs of Section 3.8 were executed on the Omic platform under the k8s_gpu profile with Nextflow 26.10.2, from the wrapper at commit da92d67f01e8571b1b65292e62192a215562bb4d around NucleoDock at commit 0b931cd23660f0b5a86654be7e96a7f37988c99e; all 139 succeeded. The repeat of Section 3.10 was one platform run of a batch workflow (run 74df79eb-2182-4254-940f-634c38476037, workflow commit ef8637e676df79be4af49df4536aaf442f3acb52, container digest as above, profile k8s; 139 of 139 tasks succeeded). The matched-input AutoDock Vina arm and the smina arm of Section 3.10 were run on a local workstation (the AutoDock Vina Python bindings; smina 2017.11.9 in its public container), not on the platform, and their prepared inputs, poses and scored tables are in the archive (review/gap-closure-2026-11-20/new_arms/). The archive carries the prepared receptor, reference ligand, SMILES input and scored top-1 pose of all 139 targets (data/

`nucleodock_io/`), so the scoring is reproducible from the archive alone and the docking is reproducible from the public NucleoDock release. NucleoDock is third-party software and is not redistributed here; it is identified by its public repository and the wrapper commit actually run. The validity results of Section 3.6 were also reproduced from a digest-pinned PoseBusters container on the Omic platform: 133 of 139 poses passed all 22 checks, with 0 of 139 poses discordant from the local run. A platform run of `rnaprep` 0.2.0 on 5 instances (run `e1cd9e1c`, revision `850856e2`) produced manifests identical to the local run; version 0.3.0, which adds the alternate-location step, has been run locally only, and no platform run of it covers the full set of 2,970 instances. The `rnaprep` image digest is recorded in the archive (`tools/rnaprep/nextflow.config`).

### S7. Literature and citation screen

Of the 689 works citing PoseBusters (Semantic Scholar 473 ∪ OpenAlex 530), seven carry a nucleic-acid term in the title and five of those also carry a validity term. A further sweep of the works citing HARIBOSS returned 57 nucleic-acid titles and none with a validity term. A literature search on 13 November 2026 found no study that measures how often deposited RNA–ligand structures fail PoseBusters checks, or that explains why they fail. The PoseBusters issue tracker had no issue mentioning nucleic acids on that date. The sweeps were run with `cite_sweep.py` and `multi_search.py`.

### S8. The false-failure rate as a function of the resolution cut-off and of the entries kept

**Table S8.** Receptor-clash failure rate at the shipped threshold of 0.75 for X-ray entries up to each resolution cut-off, with the three entry rules of Section 2.2. The 2.0 Å row with no protein and no covalent bond is the clean set of Section 3.3 (3 of 149); its interval here (0.00–4.80) is from a percentile bootstrap that pools the classes and differs in the second decimal from the 0.00–4.58 of Table 2.

| resolution cut-off, Å | entries kept | rate, % (failures of n) | 95% cluster CI, % | classes |
|---|---|---|---|---|
| 1.5 | X-ray + resolution + no protein + no covalent bond | 0.00 (0 of 25) | 0.00–0.00 | 22 |
| 1.5 | X-ray + resolution + no covalent bond (protein kept) | 3.33 (1 of 30) | 0.00–9.38 | 25 |
| 1.5 | X-ray + resolution + no protein (covalent bond kept) | 7.14 (2 of 28) | 0.00–16.67 | 23 |
| 1.8 | X-ray + resolution + no protein + no covalent bond | 1.14 (1 of 88) | 0.00–3.81 | 55 |
| 1.8 | X-ray + resolution + no covalent bond (protein kept) | 4.42 (5 of 113) | 0.94–8.41 | 71 |
| 1.8 | X-ray + resolution + no protein (covalent bond kept) | 10.91 (12 of 110) | 4.17–18.55 | 59 |
| 2.0 | X-ray + resolution + no protein + no covalent bond | 2.01 (3 of 149) | 0.00–4.79 | 83 |
| 2.0 | X-ray + resolution + no covalent | 4.21 (9 of 214) | 1.81–7.21 | 125 |

| resolution cut-off, Å | entries kept | rate, % (failures of n) | 95% cluster CI, % | classes |
|---|---|---|---|---|
| | bond (protein kept) | | | |
| 2.0 | X-ray + resolution + no protein (covalent bond kept) | 8.72 (15 of 172) | 3.86–14.55 | 88 |
| 2.2 | X-ray + resolution + no protein + no covalent bond | 1.44 (3 of 208) | 0.00–3.35 | 109 |
| 2.2 | X-ray + resolution + no covalent bond (protein kept) | 2.84 (9 of 317) | 1.17–4.88 | 184 |
| 2.2 | X-ray + resolution + no protein (covalent bond kept) | 10.29 (25 of 243) | 5.53–16.00 | 114 |
| 2.5 | X-ray + resolution + no protein + no covalent bond | 3.37 (10 of 297) | 1.22–6.03 | 145 |
| 2.5 | X-ray + resolution + no covalent bond (protein kept) | 4.50 (22 of 489) | 2.60–6.70 | 260 |
| 2.5 | X-ray + resolution + no protein (covalent bond kept) | 12.17 (42 of 345) | 7.10–17.88 | 152 |
| 3.0 | X-ray + resolution + | 5.22 (29 of 556) | 3.15–7.49 | 222 |

| resolution cut-off, Å | entries kept | rate, % (failures of n) | 95% cluster CI, % | classes |
|---|---|---|---|---|
| | no protein + no covalent bond | | | |
| 3.0 | X-ray + resolution + no covalent bond (protein kept) | 6.91 (71 of 1027) | 5.16–8.67 | 431 |
| 3.0 | X-ray + resolution + no protein (covalent bond kept) | 13.36 (84 of 629) | 9.59–17.79 | 236 |

### S9. Regression of receptor-clash failure on entry properties

**Table S9.** Logistic regression of receptor-clash failure at 0.75 (2,931 instances with a measurable contact, 527 failures, 900 equivalence classes), with standard errors clustered by class. Resolution is centered at 2.0 Å and filled with the median for the 99 solution-NMR instances, whose resolution term is absorbed by the NMR indicator.

| term | odds ratio | 95% CI | P |
|---|---|---|---|
| intercept | 0.06 | 0.04–0.08 | 2.9e-60 |
| electron microscopy | 1.13 | 0.73–1.73 | 0.59 |
| solution NMR | 0.09 | 0.02–0.39 | 0.0016 |
| protein in the entry | 1.51 | 0.99–2.31 | 0.054 |
| DNA in the entry | 1.40 | 0.87–2.26 | 0.17 |
| deposited covalent bond | 27.98 | 16.80–46.61 | 1.7e-37 |
| resolution, per Å poorer (centered at 2.0 Å) | 1.90 | 1.50–2.42 | 1e-07 |

### S10. Checks that fail the top-1 poses and the deposited ligands of the docking set

**Table S10.** Number of the 139 top-1 poses failing each PoseBusters check, with the number of those poses that are accurate (RMSD < 2.0 Å) in brackets; only checks that fail at least one pose are listed. A pose can fail several checks. The receptor-clash row is at 0.75.

| check | AutoDock Vina: poses failing (accurate) | NucleoDock: poses failing (accurate) |
|---|---|---|
| internal_steric_clash | 0 (0) | 6 (1) |
| non-aromatic_ring_non-flatness | 1 (0) | 0 (0) |
| internal_energy | 6 (0) | 13 (6) |
| minimum_distance_to_protein | 0 (0) | 53 (28) |
| minimum_distance_to_organic_cofactors | 0 (0) | 3 (1) |
| minimum_distance_to_inorganic_cofactors | 0 (0) | 6 (1) |
| volume_overlap_with_protein | 0 (0) | 13 (0) |
| ANY | 7 (0) | 63 (35) |
| receptor_clash_0.75 | 0 (0) | 53 (28) |
| receptor_clash_0.735 | 0 (0) | 51 (26) |

**Table S11.** Checks failed by the deposited ligand of the 139 docking targets.

| check | deposited ligands failing |
|---|---|
| non-aromatic_ring_non-flatness | 1 |
| minimum_distance_to_protein | 7 |
| minimum_distance_to_organic_cofactors | 3 |
| volume_overlap_with_organic_cofactors | 2 |

### S12. Success of every docking arm on the 139 targets

**Table S12.** Top-1 success, % (number of targets; 95% CI from a cluster bootstrap over the 71 equivalence classes, 10,000 resamples). Targets that were not docked count as failures. The intervals here differ in the second decimal from those of Table 4, which used 4,000 resamples.

| arm | RMSD < 2.0 Å | and valid at 0.75 | and valid at 0.735 | docked | clash failures at 0.75 | minimum relative distance |
|---|---|---|---|---|---|---|
| AutoDock Vina, deposited ligand | 30.22 (42; 21.85–39.42) | 30.22 (42; 21.85–39.42) | 30.22 (42; 21.85–39.42) | 139 | 0 | 0.753 |
| NucleoDock, SMILES (first run) | 69.06 (96; 57.25–79.14) | 43.88 (61; 33.08–56.52) | 45.32 (63; 34.19–58.33) | 139 | 53 | 0.199 |
| NucleoDock, SMILES (repeat, platform batch run) | 69.06 (96; 57.25–79.14) | 44.60 (62; 33.80–57.02) | 46.04 (64; 34.92–58.72) | 139 | 55 | 0.15 |
| AutoDock Vina, RDKit conformer of the NucleoDock SMILES | 23.02 (32; 14.46–34.95) | 23.02 (32; 14.46–34.95) | 23.02 (32; 14.46–34.95) | 139 | 0 | 0.809 |
| smina, deposited ligand (5 unreadable ligands counted as failures) | 28.78 (40; 20.42–37.75) | 28.78 (40; 20.42–37.75) | 28.78 (40; 20.42–37.75) | 134 | 0 | 0.818 |
| smina, deposited ligand (5 macrocycles prepared rigid) | 30.22 (42; 21.58–39.60) | 30.22 (42; 21.58–39.60) | 30.22 (42; 21.58–39.60) | 139 | 0 | 0.818 |

**Table S13.** Paired difference in top-1 success, NucleoDock (first run) minus each other arm.

| comparison | criterion | difference, pp (95% CI) | cluster permutation P |
|---|---|---|---|
| nucleodock minus vina_deposited | rmsd | 38.85 (26.47 to 50.00) | 0.0001 |
| nucleodock minus vina_deposited | valid_0.75 | 13.67 (1.34 to 27.52) | 0.0517 |
| nucleodock minus vina_deposited | valid_0.735 | 15.11 (2.42 to 29.20) | 0.0313 |
| nucleodock minus vina_matched_rdkit | rmsd | 46.04 (30.95 to 58.00) | 0.0001 |
| nucleodock minus vina_matched_rdkit | valid_0.75 | 20.86 (10.42 to 30.91) | 0.0011 |
| nucleodock minus vina_matched_rdkit | valid_0.735 | 22.30 (11.76 to 32.54) | 0.0004 |
| nucleodock minus smina_deposited | rmsd | 40.29 (28.93 to 50.72) | 0.0001 |
| nucleodock minus smina_deposited | valid_0.75 | 15.11 (3.08 to 28.57) | 0.0232 |
| nucleodock minus smina_deposited | valid_0.735 | 16.55 (4.23 to 30.14) | 0.0125 |
| nucleodock minus smina_deposited_rigid_macrocycles | rmsd | 38.85 (26.36 to 50.00) | 0.0001 |
| nucleodock minus smina_deposited_rigid_macrocycles | valid_0.75 | 13.67 (1.34 to 27.35) | 0.0493 |
| nucleodock minus smina_deposited_rigid_macrocycles | valid_0.735 | 15.11 (2.56 to 28.99) | 0.0273 |
| nucleodock minus nucleodock_repeat | rmsd | 0.00 (0.00 to 0.00) | 1.0000 |
| nucleodock minus nucleodock_repeat | valid_0.75 | -0.72 (-2.46 to 0.00) | 1.0000 |
| nucleodock minus nucleodock_repeat | valid_0.735 | -0.72 (-2.46 to 0.00) | 1.0000 |

### S14. The comparison inside and outside the external test set of NucleoDock

**Table S14.** NucleoDock minus another arm on the targets whose PDB entry is in the 125-complex external test set of the NucleoDock preprint, and on the others, whose use in training cannot be checked.

| subset | targets (classes) | versus | criterion | NucleoDock, % | other arm, % | difference, pp (95% CI) |
|---|---|---|---|---|---|---|
| in the 125-complex external test set | 53 (30) | vina_deposited | rmsd | 67.92 | 28.30 | 39.62 (20.93 to 55.56) |
| in the 125-complex external test set | 53 (30) | vina_deposited | valid_0.75 | 33.96 | 28.30 | 5.66 (-15.79 to 27.69) |
| in the 125-complex external test set | 53 (30) | vina_deposited | valid_0.735 | 37.74 | 28.30 | 9.43 (-12.00 to 32.14) |
| in the 125-complex external test set | 53 (30) | vina_matched_rdkit | rmsd | 67.92 | 16.98 | 50.94 (32.56 to 65.12) |
| in the 125-complex external test set | 53 (30) | vina_matched_rdkit | valid_0.75 | 33.96 | 16.98 | 16.98 (-2.50 to 36.21) |
| in the 125-complex external test set | 53 (30) | vina_matched_rdkit | valid_0.735 | 37.74 | 16.98 | 20.75 (1.89 to 40.00) |
| in the 125-complex external test set | 53 (30) | smina_deposited | rmsd | 67.92 | 24.53 | 43.40 (25.58 to 58.93) |
| in the 125-complex | 53 (30) | smina_deposited | valid_0.75 | 33.96 | 24.53 | 9.43 (-10.21 to 30.88) |

| subset | targets (classes) | versus | criterion | NucleoDock, % | other arm, % | difference, pp (95% CI) |
|---|---|---|---|---|---|---|
| external test set | | | | | | |
| in the 125-complex external test set | 53 (30) | smina_deposited | valid_0.735 | 37.74 | 24.53 | 13.21 (-6.78 to 35.00) |
| not in that set (training status unknown) | 86 (52) | vina_deposited | rmsd | 69.77 | 31.40 | 38.37 (24.36 to 51.69) |
| not in that set (training status unknown) | 86 (52) | vina_deposited | valid_0.75 | 50.00 | 31.40 | 18.60 (2.33 to 36.36) |
| not in that set (training status unknown) | 86 (52) | vina_deposited | valid_0.735 | 50.00 | 31.40 | 18.60 (2.33 to 36.36) |
| not in that set (training status unknown) | 86 (52) | vina_matched_rdkit | rmsd | 69.77 | 26.74 | 43.02 (25.61 to 58.72) |
| not in that set (training status unknown) | 86 (52) | vina_matched_rdkit | valid_0.75 | 50.00 | 26.74 | 23.26 (10.98 to 36.25) |
| not in that set (training status unknown) | 86 (52) | vina_matched_rdkit | valid_0.735 | 50.00 | 26.74 | 23.26 (10.98 to 36.25) |
| | 86 (52) | smina_deposited | rmsd | 69.77 | 31.40 | |

| subset | targets (classes) | versus | criterion | NucleoDock, % | other arm, % | difference, pp (95% CI) |
|---|---|---|---|---|---|---|
| not in that set (training status unknown) | | | | | | 38.37 (25.84 to 50.59) |
| not in that set (training status unknown) | 86 (52) | smina_deposited | valid_0.75 | 50.00 | 31.40 | 18.60 (3.09 to 35.71) |
| not in that set (training status unknown) | 86 (52) | smina_deposited | valid_0.735 | 50.00 | 31.40 | 18.60 (3.09 to 35.71) |

### S15. The receptor-clash failures of NucleoDock

The receptor-clash failures follow the NucleoDock pose and not the target. None of the 53 NucleoDock clash failures also failed the check for the AutoDock Vina pose of the same target, and 5 also failed it for the deposited ligand of that target. The most severe pose, 4YAZ with ligand 4BW, places a ligand carbon 0.655 Å from a receptor nitrogen (relative distance 0.199, 65 clashing atom pairs); the median failing pose had 2 clashing pairs. The most extreme pair was most often a ligand nitrogen with a receptor oxygen (21 poses), a ligand oxygen with a receptor nitrogen (8) or two oxygens (8), which are polar contacts at distances a hydrogen bond does not allow. The difference in ligand input is not an obvious source of the cost. The median ratio of the pose's internal energy to that of a relaxed conformer ensemble was 2.88 for AutoDock Vina, 2.30 for NucleoDock and 1.72 for the deposited ligands, so the program whose ligands were further from a relaxed geometry by this measure is the one with no clash failures. A matched-input comparison is reported in Section 3.10.

### S16. Robustness analyses in full

**Matched ligand input.** AutoDock Vina was run again from an RDKit conformer of the SMILES string that NucleoDock received (Section 2.7). It placed 32 of 139 ligands within 2.0 Å (23.02%; 95% CI 14.89–34.93), all of them valid, identically at 0.75 and 0.735 (after superposition the figure would have been 88). No pose failed the receptor-clash check (minimum relative distance 0.809), and no accurate pose failed any check. Against this arm NucleoDock led by 46.04 points on RMSD (95% cluster CI 30.95–58.00; cluster permutation $P < 0.001$) and by 20.86 points with validity required at 0.75 (10.42–30.91; $P = 0.001$) and by 22.30 at 0.735 (11.76–32.54; $P < 0.001$). The matched arm did not differ detectably from the deposited-ligand arm (−7.19 points, −15.76 to 1.64, on every criterion). The lead of NucleoDock therefore does not depend on whether AutoDock Vina starts from the deposited ligand or from the ligand geometry that NucleoDock received.

**A third program.** smina 2017.11.9 docked the deposited ligands into the same boxes. Five ligands (1ZZ5/CNY, 4QLM/2BA, 4QLN/2BA, 4YAZ/4BW and 4YB0/C2E) carry the flexible-ring pseudo-atoms that Meeko writes for macrocycles, which this smina version rejects (it reports that CG0 is not a valid AutoDock type), and were counted as failures to dock. On that basis smina placed 40 of 139 ligands within 2.0 Å (28.78%; 20.32–37.68), all of them valid. No pose failed the receptor-clash check (minimum relative distance 0.818). NucleoDock led by 40.29 points on RMSD (28.93–50.72) and by 15.11 points with validity required at 0.75 (3.08–28.57; $P = 0.023$) and by 16.55 at 0.735 (4.23–30.14; $P = 0.013$). Preparing the five macrocycles with rigid rings and docking them gave 42 accurate poses (30.22%; 21.48–39.64), all valid, a lead of 38.85 points on RMSD (26.36–50.00) and of 13.67 points with validity required (1.34–27.35; $P = 0.049$), still without a receptor-clash failure. Requiring validity cost the four AutoDock Vina and smina arms no accurate pose, against 25.18 points for NucleoDock.

**Repeat of the NucleoDock run.** The 139 NucleoDock runs were repeated with the same container digest, configuration and inputs (Section 2.7). The poses of the first run were not archived: its unaligned RMSD is taken from the repeat's poses, except for 2GDI, 2HOJ and 2HOM, whose first-run poses were read back from the platform output because their RMSD lies near the 2 Å cut (2.119, 1.857 and 1.769 Å from the first run against 2.100, 1.842 and 1.765 Å from the repeat), so the reproducibility of the RMSD is judged on the superposed value of each run. After superposition the repeat placed 125 ligands within 2.0 Å against 126 in the first run; one call changed. The superposed RMSD agreed to 0.001 Å for 119 of 139 targets, and the largest change was 0.48 Å. Without superposition the repeat placed 96 ligands within 2.0 Å (69.06%; 95% CI 57.04–79.09) and 62 with a valid pose at 0.75 (44.60%; 33.80–57.02), against 61 in the first run, so one validity-qualified call changed (a gain); the validity check alone changed on three targets (one gained, two lost). The receptor-clash check failed 55 poses in the repeat against 53 in the first run (minimum relative distance 0.150 against 0.199). The validity cost was 24.46 points in the repeat and 25.18 in the first run, and the runs differed by 0.72 points on the validity-qualified criterion (cluster CI −2.46 to 0.00), which is small against the 13.67-point lead and the 25.18-point validity cost.

**Overlap with the development data of NucleoDock.** The public repository of NucleoDock at the commit that was run ships no training, validation or test lists; it holds example inputs for four entries (2GDI, 2QWY, 5BTP and 3FU2), all four among the 139 targets and two of them (2GDI and 3FU2) in the external test set of its preprint. The preprint states that the 125 complexes of that test set were excluded from training and that 1,106 complexes were used for training, without listing them. Of the 139 targets, 53 (53 entries, 30 equivalence classes) are in the external test set, which the preprint excludes from training, and 86 (83 entries) are not, so their use in training can be neither confirmed nor excluded. On the 53 external test targets NucleoDock led AutoDock Vina by 39.62 points on RMSD (95% cluster CI 20.93–55.56) and by 5.66 points with validity required at 0.75 (−15.79 to 27.69); on the other 86 targets the corresponding differences were 38.37 points (24.36–51.69) and 18.60 points (2.33–36.36). NucleoDock's RMSD success was 67.92% on the external test targets and 69.77% on the others, which gives no sign of a contribution of training overlap, although the comparison cannot exclude one. The lead on RMSD is present in both subsets, so it does not depend on the targets that may overlap; with validity required it is clearest on the 86 targets, and its interval on the 53 includes zero.

### S17. The all-but-one rule and the PoseBusters calibration

PoseBusters did not use a 95% rule. Its paper describes borrowing a strain-energy ratio from earlier work (Wills et al., reference 29 of the manuscript) and loosening it ("Wills et al. set a ratio of 7 … We selected a less strict ratio of 100"), while its distance thresholds are set so that all, or all but one, crystal ligand passes. The all-but-one rule is therefore determined by the two most extreme contacts in a set. Applied to the 556 X-ray, no-protein, non-covalent instances at ≤ 3.0 Å, it falls to 0.413, set by contacts of 1.31 and 1.30 Å in 7ELP/GTP (2.79 Å resolution) and 6E8S/SPM (2.35 Å).
